# lncDIFF: a novel distribution-free method for differential expression analysis of long non-coding RNA

**DOI:** 10.1101/420562

**Authors:** Qian Li, Xiaoqing Yu, Ritu Chaudhary, Robbert JC Slebos, Christine H. Chung, Xuefeng Wang

## Abstract

**Motivation:** Long non-coding RNA expression data has been increasingly used in finding diagnostic and prognostic biomarkers in cancer studies. Existing differential analysis tools for RNA sequencing does not effectively accommodate low abundant genes, as commonly observed in lncRNA. We propose a novel and robust statistical method lncDIFF to detect differential expressed (DE) genes without assuming the true density on normalized counts.

**Results:** lncDIFF adopts the generalized linear model with zero-inflated exponential quasi likelihood to estimate group effect on normalized counts, and employs the likelihood ratio test to detect differential expressed genes. The proposed method and tool is suitable for data processed with standard RNA-Seq preprocessing and normalization pipelines. Simulation results illustrate that lncDIFF detects DE genes with more power and lower false discovery rate regardless of the data pattern. The analysis on a head and neck squamous cell carcinomas study also confirms that lncDIFF has better sensitivity in identifying novel lncRNA genes with relatively large fold change and prognostic value.

**Availability and Implementation:** lncDIFF is an R package available at https://github.com/qianli10000/lncDIFF.

**Supplementary Information:** Supplementary Data are available at Bioinformatics online.

## INTRODUCTION

Long noncoding RNAs (lncRNAs) are transcripts longer than 200 nucleotides with no or limited protein-coding capability. It is estimated that, in the human genome, there are at least four times more lncRNA genes than protein-coding genes [1]. Currently, there are more than 14,000 human lncRNAs annotated in GENCODE (https://www.gencodegenes.org/). Overall, lncRNA genes have fewer exons, lower abundance and are under selective constraints compared to protein-coding genes. LncRNAs are involved in diverse regulatory mechanisms and in some critical pathways. For example, they can act as scaffolds to create higher-order protein complexes, as decoys to bind sequester transcription factors, and as guides of protein-DNA interactions [2-4]. Emerging evidence suggests that lncRNA serve as essential regulators in cancer cell migration and invasion, as well as in other cancerous phenotypes [5, 6].Therefore, lncRNAs are becoming attractive potential therapeutic targets and a new class of biomarkers for the cancer prognosis and diagnosis. For example, the lncRNA PCA3 (prostate cancer antigen 3) is a FDA-approved biomarker for prostate cancer prediction. The overexpression of lncRNA HOTAIR in breast cancer patients is reported to be associated with patient survival and risk of metastasis [7]. Another important lncRNA ANRIL (CDKN2-AS1) is one of the most frequently alerted genes in human cancers and has been reported to increase cancer risks in diverse cancers.

Although a large number of lncRNAs have been identified, only a very small proportion of them have been characterized for cellular and molecular functions. Similar to protein-coding genes, the biomarker discovery of lncRNAs can start from a genome-wide differential expression (DE) analysis. One advantage of lncRNAs research in cancer is that we can leverage the large collection of previously published RNA-seq data and perform secondary analyses. Unlike the miRNAs counterparts, the expression of a large number of lncRNAs can be detected by standard RNA-seq with sufficient sequencing depth. Through downloading RNA-seq BAM files and recalling using GENCODE genomic coordinates, more than 8,000 human tumor samples across all major cancer types in The Cancer Genome Atlas (TCGA) and other published studies have been re-analyzed for the lncRNAs expression profile [8, 9]. There is a limited number of non-tumor samples sequenced for RNA-seq in TCGA. If necessary, the database such as the GTEx (http://gtexportal.org) can serve as additional tissue-specific controls, which provides over 9,600 RNA-seq samples across 51 tissues.

lncRNAs expression data have several features that pose significant challenges for the data analysis, including low abundance, large number of genes, and rough annotations. To ensure detection reliability, a common practice is to filter out lncRNA genes with low average Reads Per Kilobase per Million mapped reads (RPKM), e.g. <0.3. We recommend using the two-step filter proposed in [9]: in the first step eliminates genes with 50th-percentile RPKM =0, and in the second step only keep genes whose 90th-percentile RPKM <0.1. About two-thirds of lncRNAs are excluded after this filtering procedure.Interestingly, excess zeros or low expression values are still observed in the downsized dataset. It is well known that excess zero read counts in RNAseq data can distort model estimation and reduce power in differential expression analysis. The popular R packages DESeq2 and edgeR assume a negative binomial (i.e. over-dispersed Poisson) distribution for the count data. Methods based on zero-inflated negative binomial (ZINB) and zero-inflated GLM have been proposed to explicitly address the issue of excess zeros in RNA-seq data [10]. These methods have been recently applied to single-cell RNA-seq (scRNA-seq) data, which has high dropout rates. Since the difference in gene expression variance is biologically interesting, multiple methods have been developed to incorporate the testing of variance in the differential model. However, for biomarkers in clinical settings, genes with pronounced group contrast in mean expression level usually have more translation value. Gene wise expression variability can generate from different sources and varies widely from study to study, especially with different normalization methods. Hence, we focus on the group comparison of mean gene expression level regardless of variability in this study.

In a large-scale secondary analysis of expression data such as in lncRNA studies, only normalized data (such as RSEM or RPKM) are available [11, 12]. Certain packages such as DESeq2, however, cannot be applied because they do not accept normalized expression and zero as input. In this case, a common practice is to round continuous expression values into integers and shift it to be nonzero. Another commonly-adopted approach is using *log*_2_ (*x* + 1) transformed normalized data in R package like limma [13], i.e., assuming a log-transformed Gaussian distribution as in microarray intensity levels. The core function in limma, which basically runs a moderated t-test after an empirical Bayes correction, is more generic and more suitable for the differential expression of processed lncRNA expression data. In a very recent study, a total of 25 popular methods for testing differential expression genes were comprehensively evaluated with special emphasis on low-abundance mRNAs and lncRNAs [14]. It was observed that linear modeling with empirical Bayes moderation (implemented in limma with variance stabilizing transformation [15], voom [16] or trend), and a non-parametric method based on Wilcoxon rank sum statistic (implemented in SAMSeq) showed overall good balance of false discovery rate (FDR) and reasonable detection sensitivity. However, none of the methods compared can outperform all other tools and all tools exhibited substandard performance for lncRNAs in terms of differential testing, often with higher FDR and true positive rate (TPR) than for mRNAs. This study also concluded that accurate differential expression inference of lncRNAs requires more samples than that of mRNAs. Even methods like limma can exhibit an excess of false discoveries under specific scenarios, making these methods unreliable in practical applications.

In this paper, we present the lncDIFF, an efficient and reliable toolset based on a zero-inflated exponential quasi-likelihood strategy without the need to fully specify a parametric model. The quasi-likelihood model provides unbiased and efficient estimators even under erroneous assumptions about density. It thus provides a simple and versatile approach to model gene expression data without making strong distributional assumptions about the underlying variation, but still being compatible with existing RNA-Seq quantification and normalization tools. The flexibility in allowing for the estimation of calibration and variance parameters is especially important for lncRNAs differential analysis. The lncDIFF is thus able to integrate desirable features from the aforementioned two top-performing methods (limma and SAMSeq [14]) for lncRNA differential analysis. The lncDIFF is compared with existing tools using an extensive simulation study and real data analysis on TCGA head and neck squamous cell carcinomas (HNSC). Results suggest that lncDIFF is powerful and robust in a variety of scenarios and identifies DE lncRNA genes of low expression with more accuracy.

## METHOD

### RNA-Seq Counts Distribution Based Variation

In RNA-Seq gene expression analysis, the type of RNAs and the selected alignment, quantification and normalization tools usually have substantial impact on the distribution pattern of transcript abundance as discussed in [17], especially on the level of gene expression dispersion, i.e. the mean-variance relation. Most of the existing RNA-Seq tools, such as DESeq [18], edgeR [19], and baySeq [20] estimate gene-wise counts dispersion to perform raw counts normalization or differential expression analysis. However, this technique may not be suitable for low-abundance mRNA or lncRNA. The analysis tools such as limma [13, 21] with data transformation become superior for lncRNA instead [14]. In other words, the underlying mean-variance relation distinguishes different types of RNA-Seq counts and determines the tools for downstream analysis.

Let *X*_*gi*_ represent RNA-Seq read counts mapped to gene *g* in sample *i, g* = 1, …, *G, i* = 1, …, *N*.The existing analysis on RNA-Seq data usually assumes Negative Binomial (NB) or the Log Normal (LN) distribution for raw or normalized counts [14, 16], with mean-variance relation summarized as a quadratic form *Var*(*X*_gi_) =*c E*(*X*_gi_)^2^. The positive constant *c* is the ‘variation’ parameter, i.e., the square of coefficient of variation (CV) and depends on the density, i.e. 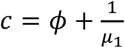 for NB and *c* = exp (σ^2^) − 1 for LN [18]. The parameters _!_, ϕ are the mean and dispersion of NB, and σ is the log standard deviation of LN, not affected by the log mean.

We use the lncRNA and mRNA data in the TCGA HNSC study to investigate the variation patterns for different types of sequencing counts. If the quantified RNA-Seq reads follow a NB distribution, the gene-wise variation or CV changes inversely with mean expression level, as revealed by the violin box plots for mRNA normalized counts in **Figure 1**. In contrast, the CV level for lncRNA normalized counts in the same study presented by **Figure 1** does not change along with gene-wise mean at 20^th^-50^th^ percentiles, similar to the LN distribution in which CV is independent of (log) mean.

**Figure 1:**
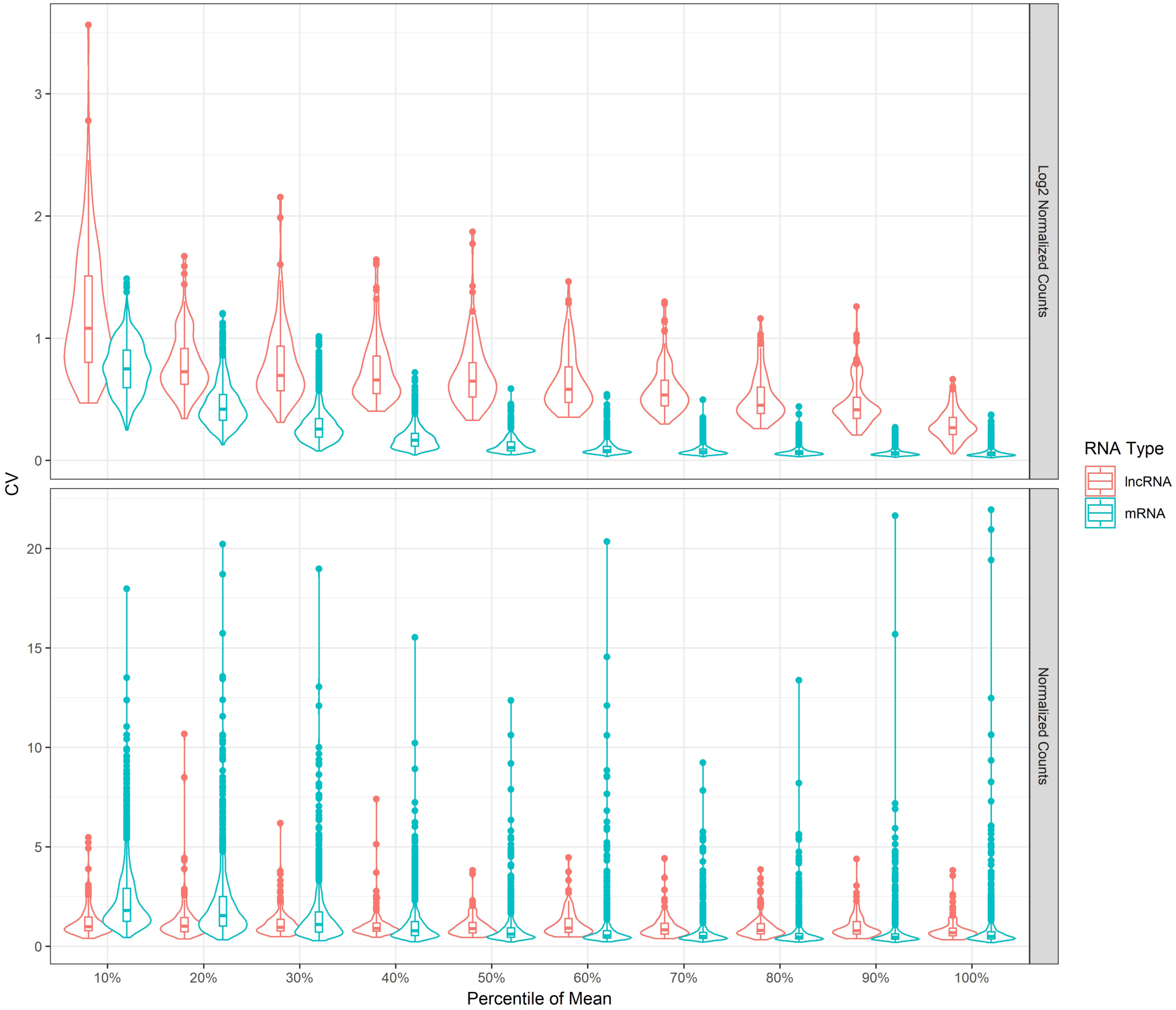
Violin and box plots for gene-wise coefficient of variation (CV) based on normalized counts of mRNA and lncRNA. Genes are divided into ten groups by the percentile of mean normalized counts. The first panel is plotted on the log2 scaled normalized counts, while the data used in the second panel is not log transformed.

For mRNA read counts, NB density is a valid assumption and provides robust estimate for the mean expression, dispersion and DE analysis. However, the scale and distribution of lncRNA counts varies across genes, some of which are extremely low and similar to LN, or have a mixture distribution of NB and LN. The differential analysis for a large number of lncRNA genes with mean expression ranging from less than 1 to over 100 can be severely biased, if one uses NB model to account for dispersion across all genes, or transformation such as log2 [22], voom [16] and variance stabilizing transformation (VST) [15] to remove the skewness for all genes. In the light of fewer statistical assumptions and parameters, it is worth to investigate the plausibility of utilizing Exponential density in RNA-Seq analysis, for which the quadratic mean-variance relation is CV = 1.

### Exponential Quasi-Likelihood

In this study, we only consider the normalized lncRNA expression data, i.e. Reads Per Kilobase per Million mapped reads (RPKM) [23] or Fragments Per Kilobase per Million mapped reads (FPKM), as the aim is to improve hypothesis testing of treatment or biological group effect on lncRNA expression regardless of the latent variation pattern. The common normalization methods, such as UQ, TMM [24, 25] are also compatible with lncDIFF, but not assessed in the simulation study and real data analysis, due to limited publically available lncRNA raw counts. We will demonstrate that the choice of normalization method does not affect the validity and accuracy of parameter estimation and DE analysis results in lncDIFF. See the last subsection of Method.

Let *Y*_*ij*_ be the lncRNA RPKM for gene *i* in sample *j*, belonging to phenotype or treatment group *k, k* = 1, …, *K*. The generalized linear model (GLM) for *Y* _*ij*_ with the Exponential family is

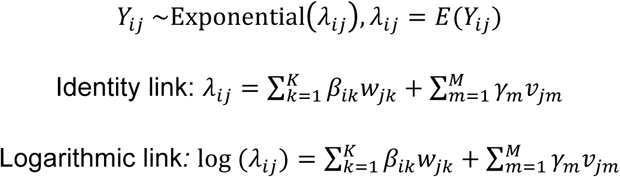

*w* _jk_ and *β* _gi_ are design matrix elements and unknown coefficients for groups, *v* _*jm*_ and *γ*_m_ are the covariates and corresponding coefficients. Since *Y* _*ji*_ has been normalized for library size, this model does not include the RNA sequencing normalization factor, although it is a common parameter in existing tools based on NB assumption [18, 19, 26, 27].

In the absence of zero expression, lncDIFF uses the Exponential GLM to lncRNA RPKM DE analysis regardless of the true density of *Y*_*ij*_ as a quasi-likelihood approach, which uses a distribution-free statistics to estimate group-wise mean RPKM, similar to the pseudo likelihood (PL) and quasi likelihood (QL) for dispersion estimate in [27]. Let *β*_*i*_ = (*β*_*i1*,…_,*β* _*ik*_) and *γ* = (*γ*_1,…_,*γ*_*m*_), for gene *i* with negligible zero occurrence (<1%), the GLM likelihood based on the exponential density 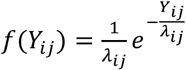 with identity or log link function is

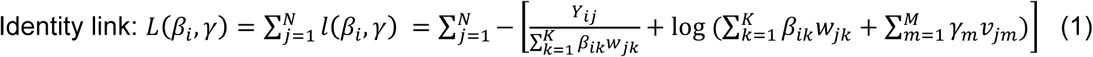

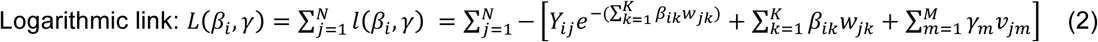

The exponential quasi likelihood estimate for mean RPKM in lncDIFF is the maximizer of *L*(*β*_*i*_, *γ*), that is 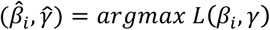. This estimate does not require prior knowledge about the statistical distribution of RPKM values, and accommodates the genes with a wide range of expression, i.e. having both extremely low (RPKM<1) and regular (RPKM>10) abundance in a large proportion of samples. The commonly adopted statistical assumptions like Poisson, NB or LN densities about RNA-Seq counts are still allowed in lncDIFF. However, the specified density does not affect the estimation of mean RPKM 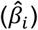 and the corresponding DE analysis results, as illustrated in the Supplementary Methods and simulation study.

### Zero-Inflated Exponential Quasi Likelihood

In lncRNA expression data, it is common to observe zero value for a gene in a non-negligible proportion (i.e., at least 1%) of samples. The excess zeroes in lncRNA RPKM cannot be addressed by integer models like Poisson and Negative Binomial (or Gamma-Poisson), since RPKM for most lncRNA genes are non-integer and fall in the range of (0, 2). Rounding decimals to integers and then applying Poisson or NB density [22, 28] or using data transformation, e.g. log2, voom, or VST [15, 16, 22] with limma [13, 21] may lead to errors in DE analysis. Therefore, we propose the zero-inflated quasi likelihood for the GLM of *Y* _*ij*_ to account for the inflation of zeros in lncRNA expression.

In order to incorporate the zero-inflated pattern, we first re-specify the RPKM for gene *i* in sample *j* by a multiplicative error model [29-31] with random error ϵ _*ij*_, that is

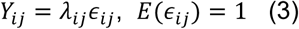

The random errors ϵ_ij_ also have the occurrence of excess zeros with a prior probability mass *P* (ϵ _gi_ = 0) = 1 - π, *P* ϵ_*ij*_ > 0 = π, and a continuous density at positive value with *E*(ϵ _ij_ |*Y* _ij_ > 0) = γ, similar to [30, 32, 33]. If the positive RPKM *Y* _ij_ |*Y* _ij_ > 0 follows the Exponential distribution (so does ϵ _ij_ *Y* _ij_ > 0, then the density functions for *Y* _ij_ including zero occurrence is

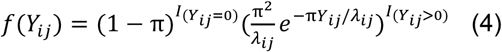

Equation (4) is derived in the Supplementary Methods. See supplementary data files. Similar to the aforementioned Exponential quasi-likelihood for GLM, lncDIFF applies the zero-inflated density in equation (5) to GLM as a quasi-likelihood approach to perform DE analysis of zero-inflated lncRNA expression. The corresponding quasi-likelihood function is

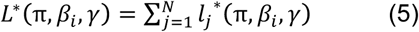

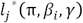 is defined according to the selected link function as

Identity link:

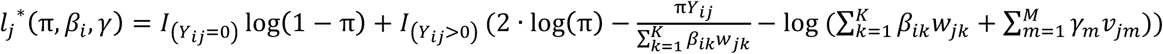

Logarithmic link: 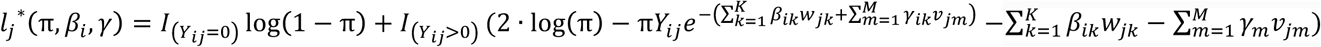

The zero-inflated quasi-maximum likelihood (ZI-QML) estimate for group-wise mean RPKM is the maximizer of *L*\* (π, β_*i*_, γ) in equation (6), that is

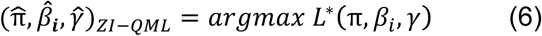

It is worthwhile to note that the likelihood function *L*\* (π, β_*i*_, γ) in equation (5) reduces to equations (1) and (2) if the proportion of zero expression is negligible, i.e. no more than 1%.

### Likelihood Ratio Test

For differential analysis in lncDIFF, we apply the Likelihood Ratio Test (LRT) to the zero-inflated exponential likelihood function *L*\* (π, β_*i*_, γ) to test hypothesis: *H*_0_: *β*_*i*_ = *β*_*null*_ vs *H*_1_: *β*_*i*_ = *β* _*full*_, where *β*_*null*_ is the design matrix coefficients with some equal to zero and *β* _*full*_ is the coefficients without zero. The test statistic of LRT is *D* = −2*L*\* (*β*_*null*_) + 2*L*\*(*β*_*null*_) with *β* _*null*_ and *β* _full_ being the design matrix coefficients for null and alternative models. Let *m* _*null*_ and *m* _*full*_ be the number of distinct coefficients in *β* _*null*_ and *β* _*full*_. Test statistic *D* asymptotically follows *χ*^2^ distribution with degrees of freedom *m* _*full*_ - *m* _*null*_. The p-values from LRT are adjusted for multiple testing using the procedure of Benjamin and Hochberg false discovery rate [34]. The choice of link function does not affect the power of LRT, as illustrated by simulation study.

We also provide empirical distribution of LRT statistics *D* to compute the p-values for DE analysis, similar to [28]. The empirical distribution of statistics *D* per gene can be generated by randomly shuffling the samples into *K* groups for *P* times and then calculate the LRT statistics for each permutation, that is *D*_1_, …, *D*_*p*_. Let the test statistics for the true groups be *D*_0_, then the empirical p-value is 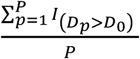,and can be adjusted by Benjamin and Hochberg procedure. We implemented the LRT for lncRNA DE analysis based on ZI-QML with observed and empirical p-values.

### lncDIFF on Other Normalization Methods

lncDIFF adopts the estimator 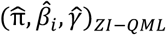 in equation (6) to estimate the mean gene expression level, based on a likelihood function that captures zero-inflation pattern without assuming the true density of non-zero RPKM. We can theoretically prove that this estimate is asymptotically unbiased, i.e., 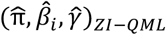 converges to the true value of (π, *β*_*i*_, *γ*) as sample size increases.

According to [35, 36], 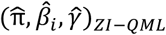 is asymptotically unbiased as long as *L*\* (π, β_*i*_, γ) converges to 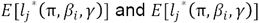 is uniquely maximized at the true value, i.e. π_0_, *β*_*i*0_, *γ*_0_. Suppose *β*_*i*0_ = *β*_*i10*_, …, *β* _*ik0*_), *γ*_0_ = (*γ* _10,…_,*γ*_*m0*_), for identity link function, the true expectation of *Y* _*ij*_ is 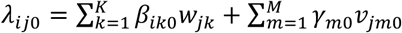. By law of large numbers, it is not hard to show that *L*\* (π, β_*i*_, γ) converges to 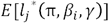, where

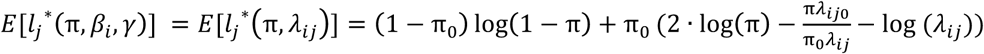

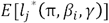 being uniquely maximized at (π_0_, *β*_*i0*_, *γ*_0_) is demonstrated by maximizing the term 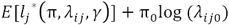, which does not depend on the distribution assumption about non-zero lncRNA RPKM. Detailed proof for unbiased estimate in either link function is elaborated in the Supplementary Methods. The use of Exponential family quasi likelihood in lncDIFF guarantees the accuracy of mean expression estimate under unknown distribution of RPKM *Y* _*ij*_. Thus, lncDIFF mean expression estimation on other normalized counts, (i.e., normalized by TMM or UQ) is also asymptotically unbiased, as long as the normalized counts are non-negative.

In order to illustrate normalization method having no impact on lncDIFF performance, we simply applied lncDIFF DE analysis to three different types of normalized counts (i.e., FPKM, TMM and UQ) of low abundance mRNA in TCGA HNSC tumor-normal samples (N=546). The low abundance genes are selected with mean FPKM in the range of (0.3, 2) and no more than 20% zero expression, similar to the majority of lncRNA genes. The Pearson correlation of log10 adjusted p-values between the three normalization methods are FPKM vs TMM 0.82, FPKM vs UQ 0.92, TMM vs UQ 0.96, implying similar DE analysis results. Therefore, we only use RPKM of lncRNA in TCGA HNSC study to illustrate the application and performance of lncDIFF in Results.

In addition to TMM and UQ, the distribution-free parameter estimation and LRT in lncDIFF are also compatible with model-based RNA-Seq quantification and normalization tools, such as RSEM [37], baySeq [20], and QuasiSeq [38]. Hence, the lncDIFF DE analysis can be incorporated into existing RNA-Seq quantification and normalization pipeline, regardless of the models employed in the preprocessing tools.

## RESULTS

### Simulation study to assess lncDIFF performance

We conducted a comprehensive simulation study to demonstrate the performance of ZI-QML with observed p-value of LRT and compare to existing common tools DESeq2, edgeR and limma (with log transformation). We rounded decimals to integers as input for DESeq2 and selected the quasi-likelihood estimation method in edgeR. The testing methods for DE genes were LRT in DESeq2 and edgeR, F-test in limma. We considered NB and LN as true densities for data sampling, and used the gene-wise estimate for dispersion or log variance from a real lncRNA RPKM dataset to determine the values of ϕ (NB) and σ2 (LN) in data generating functions. Based on the dispersion and log variance estimate for the data in TCGA head and neck squamous cell carcinomas (HNSC) study [39], we adopted ϕ = 1, *2*, 10, 20,σ^2^ = 0.01, 0.25, 1, *2*.25, and then used fixed ϕ, σ^2^ values to generate RPKM of each genes across all samples in the same simulation scenario. Each scenario is defined by the unique gene-wise nonzero proportion π = 0.5, 0.*7*, 0.9, 1, sampling density function (NB or LN) and value of ϕ, σ2, with sample size varying at N=100, 200, 300.

In order to generate data similar to lncRNA RPKM, we first obtain binary outcomes (0-1) for all samples in one scenario from the Bernoulli sampling, and then replace the 1’s by positive values generated by NB or LN densities. It should be noted that in our model with identity link function, the expectation of non-zero RPKM per gene per sample is 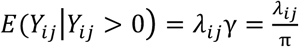. Hence, the non-zero RPKM for gene *i* in group *k* are randomly generated from NB or LN densities with mean at 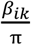, where *β*_*ik*_ being the mean of gene *i* (including zero expression) in group *k*. The HNSC study includes 40 pairs of matched normal-tumor tissues. We use the 40 normal samples to calculate the mean RPKM as baseline group parameter *β*_*i1*_ in simulation. Similar to the common filtering criteria in existing lncRNA analysis, we remove the genes in the real data with mean RPKM <0.3 [40, 41] and zero expression in more than half of the samples, reducing to 1100 genes for simulation.

In the simulation study, we only considered two-group comparison to illustrate the contrast between different methods. RPKM of the first group is randomly-generated by the specified density function and the baseline parameter, while the second group has the mean parameter of the baseline times a shift, i.e., the tumor/normal fold change in TCGA HNSC data. We manually set the shift between two simulated groups at 1 if the absolute log2 fold change (LFC) for the corresponding gene is less than 0.5. Simulated genes with between-groups shift at 1 are the null genes and the remaining are DE genes. For each simulated scenario, we generate 100 replicates to assess the performance of different methods by the mean of Type I error, false discovery rate (FDR), true positive rate (TPR), and area under the curve (AUC) of receiver operating characteristics (ROC) with FDR threshold 0.05.

We order the scenarios by the level of variance (with 1-4 representing the smallest to the largest and determined by dispersion or log variance), proportion of nonzero expression, and sample size to investigate the impact of parameters on performance metrics. **Figure 2** and **Supplementary Figures S1-S3** present the AUC, FDR, TPR and Type I error (or false positive rate) of all scenarios, illustrating that ZI-QML outperforms the other methods, especially for scenarios with LN density. AUC for all methods in **Figure 2** decrease as the gene-wise variation increases, and it also shows that ZI-QML’s performance is close to the optimal method (DESeq2) for NB density. The change of AUC across different sample sizes implies that adding more samples improves the performance of ZI-QML and DESeq2, but does not have impact on edgeR and limma. Furthermore, the AUC of ZI-QML in NB density is equivalent to or slightly larger than that of DESeq2 at sample size N=400. According to AUC and TPR, the outperformance of DESeq2 compared to lncDIFF for NB density is not as pronounced as the outperformance of ZI-QML compared to DESeq2 for LN density. On the other hand, the FDR and Type I error show that lncDIFF has similar performance of DESeq2 in most scenarios regardless of density and greatly outperforms the other two methods, although lncDIFF in large-variance LN scenarios presents performance close to edgeR and limma. In summary, lncDIFF is the optimal method for DE analysis of lncRNA RPKM with different distributions, and DESeq2 is an ideal tool if the non-zero abundance is relatively high and follow NB density.

**Figure 2:**
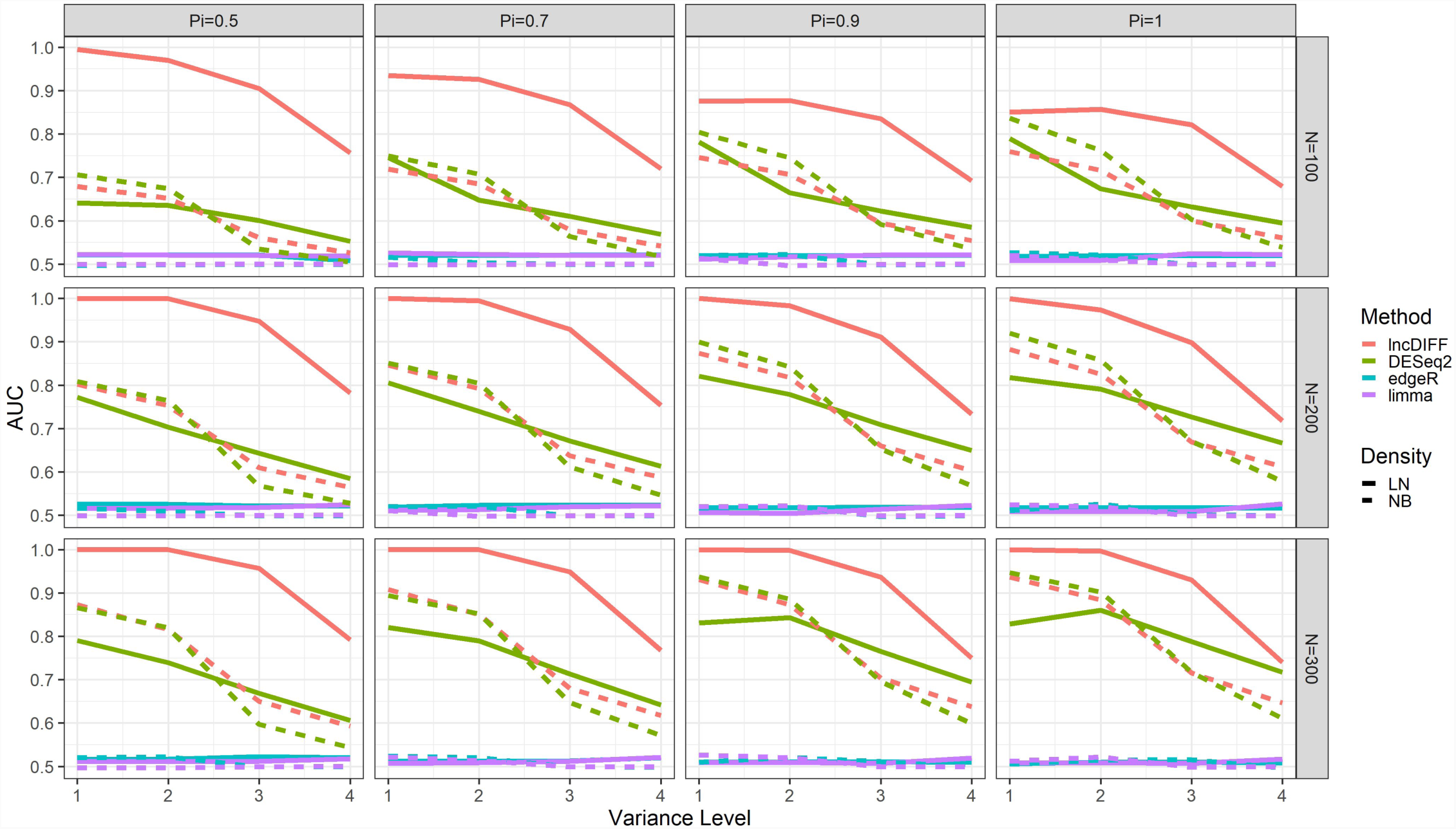
AUC of ROC curve for DE analysis on simulated data. Scenarios are in the order of true density, proportion of non-zero expression values, variance level. The labels ‘Variance1’-‘Variance4’represent gene-wise variance levels from the smallest to the largest.

### Application of lncDIFF to TCGA HNSC Data

We first employed the same methods to perform DE analysis on the TCGA HNSC lncRNA data for matched tumor and normal samples. The Venn diagram in **Figure 3 (A)** shows the overlap and difference of the DE genes identified by four methods. In real data analysis, the proportion of genes with absolute log2 fold change >0.5 and identified as DE is an alternative metric for the true positive rate, while the false positive rate (FPR) can be approximated by the proportion of genes with absolute LFC less than 0.5, 1, 1.5 but identified as DE. The significance threshold for tumor vs normal is set at FDR<0.05. We listed the alternative TPR and FPR in **Figure 3 (B)**, and presented the contrast between lncDIFF and the other methods by boxplots in **Figure 3 (C)-(E)**, with each panel showing the tumor vs normal group effect on the lncDIFF positive genes identified as negative by other methods. We only include the genes with upregulation for normal tissues and LFC>0.5 in the boxplots. The results in **Figure 3 (B)** confirms that lncDIFF provides ideal power or alternative TPR (75%) in DE analysis for LFC<0.5, with approximated FPR below 5%. The other methods either have TPR no more than 30% or generates false positives with an approximate probability 44%. The boxplots in **Figure 3 (C)-(E)** reveal that the group contrast on DE genes identified only by lncDIFF is larger than that identified only by the other methods. This also implies that lncDIFF is less likely to ‘miss’ the DE genes with large group contrast.

**Figure 3:**
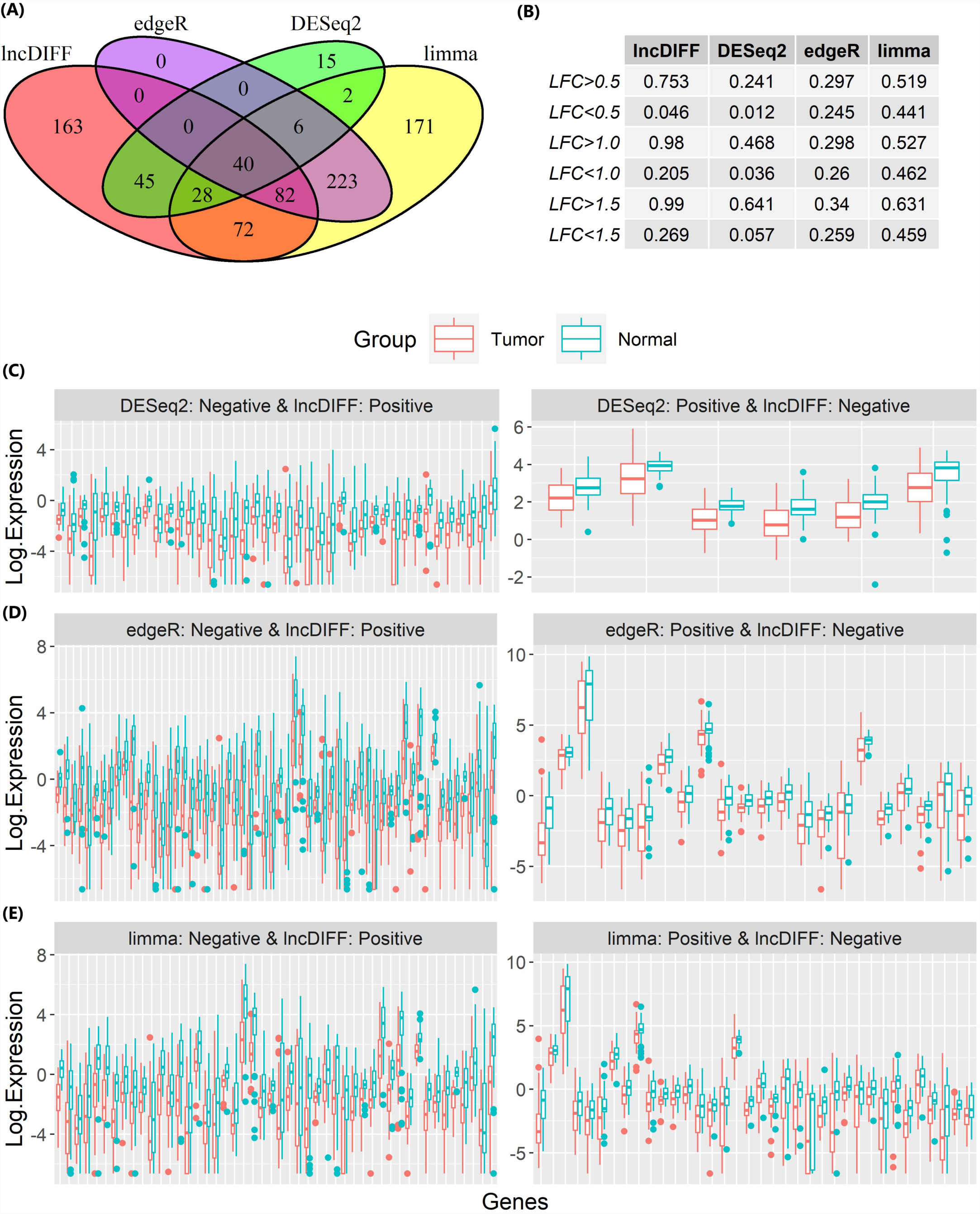
Performance of lncDIFF, DESeq2, edgeR and limma on TCGA HNSC matched tumor-normal samples. (A) is the Venn diagram for DE genes identified by each method. (B) lists the proportion of genes with LFC greater or less than 0.5, 1.0, 1.5 being identified as DE by each method. (C)-(E) are the boxplots of log2 RPKM per gene for tumor vs normal. The genes in (C)-(E) are upregulated in normal tissue and LFC>0.5.

We also applied the same analysis to the unpaired tumor (N=426) vs normal (N=40) samples in the TCGA HNSC study by lncDIFF, and compared the top significant genes in the paired and unpaired DE analysis results, listed in **Table 1**. There are 11 overlapped genes in the top-20 significant gene list of paired and unpaired analysis, some of which are associated with overall survival time. For each the overlapped significant genes, we divided the 426 HNSC tumor samples into two groups by the median of RPKM per DE gene, and then apply Cox Proportional Hazard model to survival association analysis. The Kaplan-Meier curves and the log-rank test p-values reveal marginal or significant association between genes *ERVH48-1, HCG22, LINC00668, LINC02582* and the overall survival months, illustrated by **Figure 4**. For the same set of HNSC tumor samples, we also used the mRNA RSEM normalized counts to select 20 mRNA genes highly correlated with the 11 tumor-normal DE lncRNA genes by Spearman correlation, listed in the **Supplementary Table 1**.

**Figure 4:**
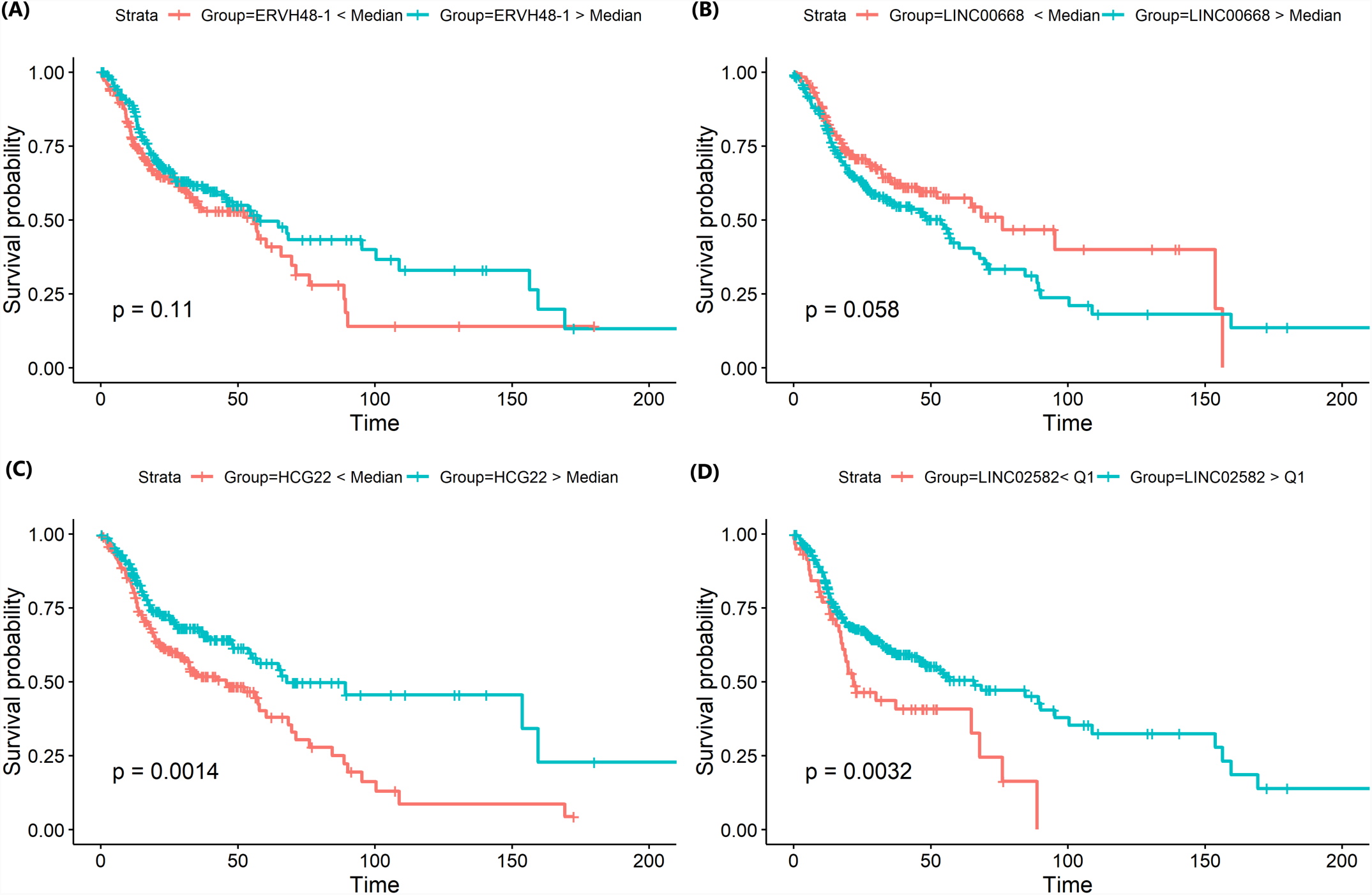
Survival time association with DE genes identified in both paired and unpaired TCGA HNSC tumor vs normal analysis. The 426 tumor samples are divided into two groups by the median of RPKM per gene. (A)-(D) are the Kaplan-Meier survival curves for genes *ERVH48-1, LINC00668, HCG22, LINC02582 individually.*

**Table 1:** lncDIFF output for the top 20 significant genes for paired and unpaired tumor vs normal differential analysis for TCGA HNSC study. The overlap of genes are in bold. Likelihood Ratio Test statistics, p-value and FDR are output from lncDIFF.

| Paired Tumor vs Normal |  |  |  |  | Unpaired Tumor vs Normal |  |  |  |  |
| --- | --- | --- | --- | --- | --- | --- | --- | --- | --- |
| Gene | Ensembl ID | Log2 Fold Change | Statistics | FDR | Gene | Ensembl ID | Log2 Fold Change | Statistics | FDR |
| **RVH48-1** | **ENSG00000233056.1** | 0.415 | 211.767 | 7.48E-45 | **HCG22** | **ENSG00000228789.2** | -2.979 | 674.029 | 1.76E-145 |
| **VC02487** | **ENSG00000203688.4** | -3.747 | 200.441 | 1.11E-42 | **LINC02487** | **ENSG00000203688.4** | -3.470 | 625.994 | 2.46E-135 |
| **HCG22** | **ENSG00000228789.2** | -3.138 | 151.425 | 3.73E-32 | MYHAS | ENSG00000272975.1 | -0.935 | 324.216 | 7.70E-70 |
| **VC00668** | **ENSG00000265933.1** | 2.189 | 148.534 | 1.20E-31 | LINC01405 | ENSG00000185847.3 | -1.366 | 276.487 | 1.45E-59 |
| **IC02582** | **ENSG00000261780.2** | 1.027 | 144.294 | 8.10E-31 | FALEC | ENSG00000228126.1 | -1.721 | 252.647 | 1.82E-54 |
| **VC00941** | **ENSG00000235884.2** | 2.450 | 138.020 | 1.59E-29 | TMEM238L | ENSG00000263429.3 | -2.250 | 235.559 | 8.06E-51 |
| VC00942 | ENSG00000249628.2 | 1.105 | 128.195 | 1.92E-27 | AC005392.2 | ENSG00000231412.2 | -2.342 | 198.936 | 6.73E-43 |
| VC01234 | ENSG00000249550.2 | 1.755 | 121.173 | 5.79E-26 | AC140479.4 | ENSG00000261760.2 | -1.471 | 188.314 | 1.23E-40 |
| VC02154 | ENSG00000235385.1 | 2.099 | 120.529 | 7.12E-26 | **ERVH48-1** | **ENSG00000233056.1** | 0.444 | 185.507 | 4.47E-40 |
| **134312.5** | **ENSG00000261327.3** | 2.064 | 115.828 | 6.85E-25 | AC091563.1 | ENSG00000254343.2 | -2.185 | 174.352 | 1.10E-37 |
| **365181.2** | **ENSG00000272068.1** | 1.191 | 111.605 | 5.24E-24 | **LINC02582** | **ENSG00000261780.2** | 1.009 | 161.008 | 8.19E-35 |
| **UXAP9** | **ENSG00000225210.5** | 2.868 | 110.895 | 6.87E-24 | **LINC00668** | **ENSG00000265933.1** | 1.626 | 154.270 | 2.23E-33 |
| **UXAP8** | **ENSG00000206195.6** | 2.422 | 105.798 | 8.30E-23 | ACBD3-AS1 | ENSG00000234478.1 | -1.733 | 150.782 | 1.08E-32 |
| IFTA1P | ENSG00000225383.2 | 1.676 | 103.239 | 2.80E-22 | **LINC00941** | **ENSG00000235884.2** | 2.090 | 150.692 | 1.08E-32 |
| 010343.3 | ENSG00000250697.1 | 1.838 | 101.397 | 6.63E-22 | **AC134312.5** | **ENSG00000261327.3** | 2.146 | 150.725 | 1.08E-32 |
| FN1-AS1 | ENSG00000236081.1 | 1.590 | 101.238 | 6.74E-22 | **DUXAP9** | **ENSG00000225210.5** | 2.711 | 141.890 | 8.49E-31 |
| VC00520 | ENSG00000258791.3 | 1.570 | 98.359 | 2.71E-21 | ABHD11 | ENSG00000225969.1 | -1.730 | 140.148 | 1.92E-30 |
| **134312.2** | **ENSG00000260162.2** | 1.912 | 98.157 | 2.84E-21 | **AL365181.2** | **ENSG00000272068.1** | 1.008 | 138.932 | 3.35E-30 |
| 114956.2 | ENSG00000248554.1 | 3.038 | 96.948 | 4.95E-21 | **DUXAP8** | **ENSG00000206195.6** | 2.230 | 134.501 | 2.95E-29 |
| CASC9 | ENSG00000249395.2 | 4.019 | 91.046 | 9.28E-20 | **AC134312.2** | **ENSG00000260162.2** | 1.982 | 129.028 | 4.42E-28 |

Secondly, we used 72 TCGA HNSC tumor samples with valid Human Papillomavirus (HPV) status (i.e. positive vs negative) to compare DE analysis results by different methods. The FDR threshold is set at <0.1 for this analysis, since the contrast between HPV positive and negative is less pronounced compared to tumor vs normal. The Venn diagram and table in **Supplementary Figure S4 (A)-(B)** show the overlap of DE genes between different methods along with the approximate power and FPR. lncDIFF still provides more power with FPR controlled at 0.02, while the other methods have DE analysis power close to either zero or FPR. **Supplementary Figure S4 (C)** are the PCA plots generated by the top 200 significant genes in terms of the FDR of each method. Based on the distance between clusters, lncDIFF top significant genes differentiate the HPV status better than those identified by the other methods do.

## DISCUSSION

lncDIFF is an efficient and powerful differential analysis tool for lncRNA RPKM or FPKM. The distribution of lncRNA RPKM is different from that of mRNA, as some genes in lncRNA may have low or even zero expression for a subset of samples, but also have normal expression for the remaining samples. Existing RNA sequencing analysis tools based on a unique density assumption ignore such characteristic and does not take excess of zeros into account. For example, DESeq2 does not allow zero counts and decimals as input data; hence, RPKM must be rounded and transformed to nonzero integers, which reduces the variation of low abundance genes across samples. Although edgeR handles non-integer RNA-Seq expression data and allows zero values, the computation for group effect depends on the estimate of gene-wise dispersion, which can be severely biased for a gene having both normal and low expression occurrence.

The Exponential likelihood function used in lncDIFF is not derived from the true density of lncRNA RPKM, but the group effect estimate based on that is valid and asymptotically unbiased, as demonstrated by the proof in Supplementary Methods. It is worth to note that this result does not hold for Poisson, NB, or LN likelihood function if the gene expression density is incorrectly assumed, since the demonstration is based on the unique structure of Exponential density function and not applicable to other distribution families. The choice of link function does not have any impact on the group effect estimate and LRT results as shown by **Table 2**, but the log link function can avoid NA values produced in numerical optimization of the likelihood function.

**Table 2:** Group effect estimates and likelihood ratio test results of TCGA HNSC tumor vs normal with logarithmic and identity link functions in lncDIFF.

| Genes Ensembl ID | Logarithmic link function |  |  | p-value | FDR |
| --- | --- | --- | --- | --- | --- |
| | $\exp(\beta_1)$<br>(tumor) | $\exp(\beta_2 - \beta_1)$<br>(contrast) | $\exp(\beta_2)$<br>(normal) | | |
| ENSG00000005206.12 | 0.247 | 0.811 | 0.200 | 0.348 | 0.528 |
| ENSG00000100181.17 | 0.737 | 0.993 | 0.732 | 0.974 | 0.982 |
| ENSG00000126005.11 | 7.161 | 1.263 | 9.043 | 0.297 | 0.474 |
| ENSG00000130600.11 | 181.885 | 1.571 | 285.661 | 0.044 | 0.115 |
| ENSG00000131484.3 | 0.362 | 1.044 | 0.378 | 0.846 | 0.916 |
| Genes Ensembl ID | Identity link function |  |  | p-value | FDR |
| | $\beta_1$<br>(tumor) | $\beta_2 - \beta_1$<br>(contrast) | $\beta_2$<br>(normal) | | |
| ENSG00000005206.12 | 0.247 | -0.047 | 0.200 | 0.348 | 0.528 |
| ENSG00000100181.17 | 0.737 | -0.005 | 0.732 | 0.974 | 0.982 |
| ENSG00000126005.11 | 7.160 | 1.887 | 9.047 | 0.297 | 0.474 |
| ENSG00000130600.11 | 181.852 | 103.833 | 285.684 | 0.044 | 0.115 |
| ENSG00000131484.3 | 0.362 | 0.016 | 0.378 | 0.846 | 0.916 |

The distribution of p-values from lncDIFF is also investigated and compared with the other methods in TCGA HNSC tumor vs normal analysis, using simulated p-values from sample permutation. We randomly selected three genes with different RPKM density patterns to generate the null p-values and then visualized the p-values distribution via QQ plots in **Figure 5**. **Figure 5** (B) shows that the p-values of lncDIFF and DESeq2 are similar and close to the expected distribution, while edgeR and limma tend to provide a large proportion of small p-values (<0.1). The histogram and density plot of RPKM presented in **Figure 5** (A) imply that the null p-values of lncDIFF and DESeq2 follow the uniform distribution for normally or highly expressed lncRNA genes (ENSG00000130600.11), and may deviate from the theoretical distribution for low abundance genes (ENSG00000152931.7, ENSG00000153363.8).

**Figure 5:**
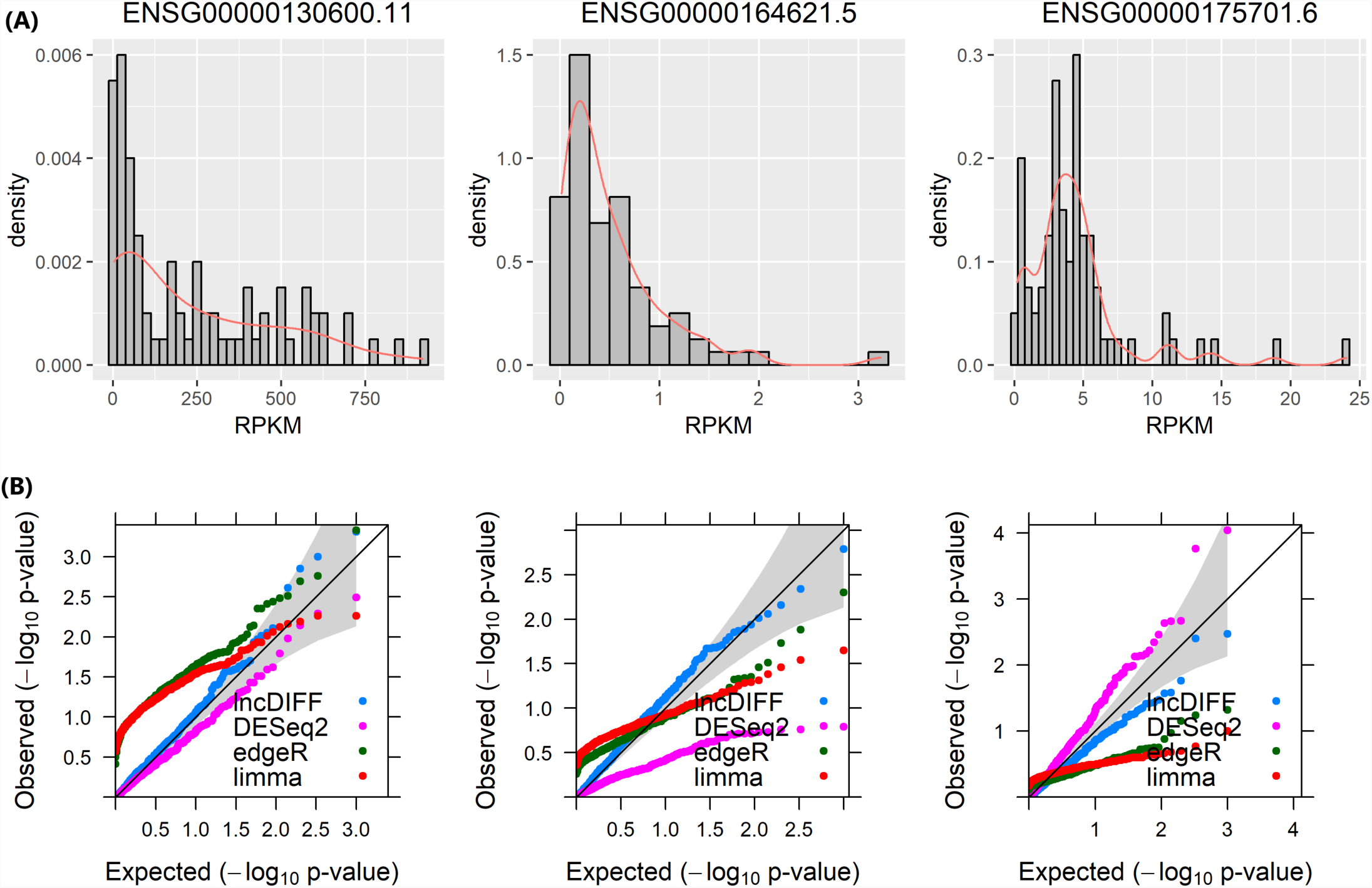
QQ plots of simulated null p-values for genes with different RPKM distributions in TCGA HNSC matched samples. (A) presents the histogram and density plot of RPKM for each genes. (B) shows the corresponding QQ plot of null p-values simulated by shuffling the samples.

We implemented ZI-QML and LRT with either link function in lncDIFF, along with an option of simulated p-values and FDR generated from permutations. This package allows the input expression matrix to be either continuous or discrete and requires group or phenotype factor provided in design matrix format. This package does not contain raw counts normalization functions, but is compatible with non-negative normalized counts from existing RNA-Seq analysis tools. The group effect estimation is implemented in R function ZIQML.fit, separated from likelihood ratio testing included in function ZIQML.LRT.

We also illustrated the computation efficiency of lncDIFF by running on the TCGA HNSC matched tumor-normal samples with ∽1130 filtered genes. The processing time (in seconds) of this biological data analysis by lncDIFF, DESeq2, edgeR and limma are 3.17, 4.31, 3.37 and 0.02, respectively. If the option of simulated p-value is enabled, the running time of lncDIFF on this real dataset is prolonged to 267.86 seconds for default 100 permutations, but the correlation between observed and simulated p-values or FDR’s is around 0.9. In future study, we will extend the lncDIFF to account for technical excess zeros from biological zeros [10], as well as apply it to lncRNA counts normalized by other methods, such as UQ and TMM.

## CONCLUSION

lncDIFF is a novel method utilizing GLM with a distribution free estimator and LRT in differential analysis of lncRNA normalized counts. This is an efficient DE analysis method, being compatible with various RNA-Seq quantification and normalization tools.

## ACKNOWLEDGEMENT

This work was supported in part by the Environmental Determinants of Diabetes in the Young (TEDDY) study, funded by the National Institute of Diabetes and Digestive and Kidney Diseases (NIDDK).

## FUNDING

This work was supported in part by Institutional Research Grant number 14-189-19 from the American Cancer Society, and the National Cancer Institute, part of the National Institutes of Health under grant number [P50 CA168536], Moffitt Skin Cancer SPORE.

## CONFLICT OF INTEREST

None declared

## Supplementary methods

### 1. Zero-Inflated Exponential density for RPKM *Y*_*ij*_

For the multiplicative error model specified in equation (3), if the positive random error ϵ _*ij*_|*Y* _*ij*_ > 0 follows a distribution described by an Exponential density function 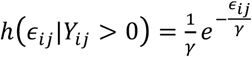 then the distribution of ϵ _*ij*_ including zero occurrence is

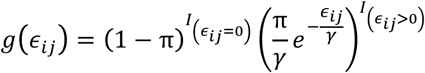

with *E*(ϵ_*ij*_) = πγ. According to the unit mean assumption *E*(ϵ _*ij*_) = 1 in equation (3), we have 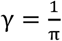. Thatis,

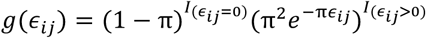

Since ϵ _*ij*_ = *Y* _*ij*_/*λ* _*ij*_, the semi-continuous distribution for *Y* _*ij*_ can be derived by

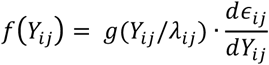

That is, 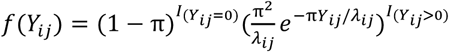

### 2. ZI-QML estimate 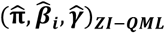 is asymptotically unbiased

Proof: According to [1, 2], 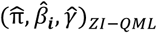 is a consistent estimator if *L*\* (π, β_*i*_, γ) converges almost surely to *E*[*l*_*j*_* (*π, β*_*i*_, *γ*)] and *E*[*l*_*j*_* (*π, β*_*i*_, *γ*)] is uniquely maximized at the true mean of RPKM, i.e. *β*_*i*0_. Suppose the true value of *β*_*i*_, *γ* are *β*_i0_ = *β*_i10_, …, *β* _ik0_, *γ*_0_ = (*γ*_*i*10,…_,*γ*_*m*0_) and the true value of π is π_0_.

**A**. Identity link function: 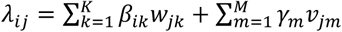

The true expectation of *Y* _*ij*_ is 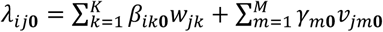 Since *E*(*y*_*ij*_ |*y*_*ij*_ > 0) = *λ*_*ij*_/π with true value *λ* _*ij*0_/π_0_, it is not hard to show that *λ*_*ij0*_/π is

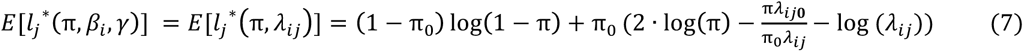

which is a finite function. By law of large numbers, *L*\* (π, β_*i*_, γ)-the sample mean of *l*\* (π, β_*i*_, γ) -converges almost surely to *E*[*l*_*j*_* (*π, β*_*i*_, *γ*)]. Next, we need to demonstrate *E*[*l*_*j*_* (*π, β*_*i*_, *γ*)] being uniquely maximized at (π_0_, *β*_i0_, *γ*_0_).

We consider the maximizer of *A*(π, *λ*_*ij*_) *E*[*l*_*j*_* (*π, β*_*i*_, *γ*)] instead of *E*[*l*_*j*_* (*π, β*_*i*_].

That is

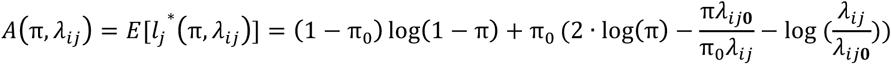

Let 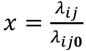, then

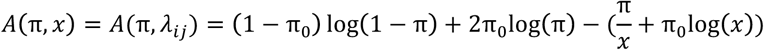

The first gradient of *A* (π, *x*) is

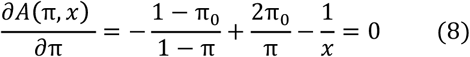

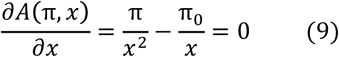

The solution to equations (8) and (9) is (π, *x*) = (π_0_, 1). The second gradient gives the Hessian matrix

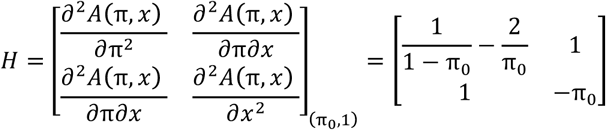

Since 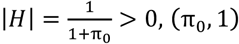 is the unique maximizer of *A* (π, *x*). Hence, (π_!_, *λ* _*ij*0)_ is the unique solution maximizing *E*[*l*_*j*_* (*π, β*_*i*_, *γ*)].

Lastly, we need to validate that *λ* _*ij*_ = *λ* _*ij* 0_ implies (*β*_*i*_, *γ*) = (*β*_*i*0_, *γ*_0_). According to definitions, the design matrix, (*β*_!_, *γ*) and *λ* _gi_ can be written as

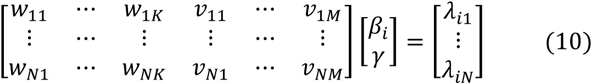

When *λ* _*ij*_ = *λ* _*ij* 0_, equation (10) becomes

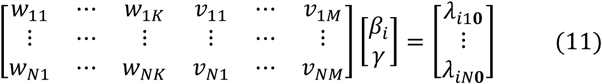

and (*β*_*i*0_, *γ*_0_) is a solution to equation (11). Since the design matrix is of full rank, the solution to equation (9) is unique. Therefore, *λ* _*ij*_ = *λ* _*ij*0_ implies (*β*_*i*_, *γ*) = (*β*_i0_, *γ*_0_), and (π_0_, *β*_i0_, *γ*_0_) is the unique maximizer of *E*[*l*_*j*_* (*π, β*_*i*_, *γ*)]. The estimator 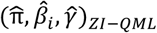 derived from *l*\* (π, β_*i*_, γ) is asymptotically consistent.

**B**. Logarithmic link function: 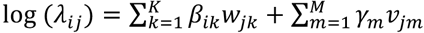

The true expectation of *Y* _*ij*_ 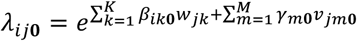Similarly, we can derive *E*\* (π, β_*i*_, γ) as equation (7) and *L*\* (π, β_*i*_, γ) converges almost surely to *E*\* (π, β_*i*_, γ), which is uniquely maximized at (π_!_, *λ*_ij0_). Similar to the above proof, 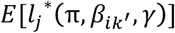 is uniquely maximized at (π_0_, *β*_i0_, *γ*_0_). Hence, consistency still holds for log link function.

